# Constrained Geometric Approach for Robust Local Optic Axis Estimation with Single Input Polarization Sensitive Optical Coherence Tomography

**DOI:** 10.64898/2026.11.13.766340

**Authors:** Jiachen Wan, Christopher R. Clickner, Maxina Sheft, Georgia L. Jones, Hui Wang

## Abstract

Polarization-sensitive optical coherence tomography (PSOCT) enables label-free, depth-resolved microstructure mapping in biological tissues, making it a powerful tool for studying myelinated fiber structures in the brain. Conventional single-input PSOCT measures cumulative retardance and apparent optic axis, which become inaccurate when fibers cross over depth. In single-input PSOCT systems, local optic axis estimation is challenging due to high noise levels, limiting the reliability of depth-resolved orientation estimation. In this work, we present a robust framework for local optic axis estimation in single-input PSOCT that incorporates physical constraints and polarization super-pixels. Polarization super-pixels serve as a preprocessing step to cluster spatially connected pixels with similar polarization states, increasing the effective signal-to-noise ratio (SNR) and reducing instability in recursive matrix inversion. In addition, a physical constraint based on the mirror state of the input polarization is enforced during geometric fitting, significantly improving robustness and accuracy under noisy and low-SNR conditions. Experimental validation on ex vivo human brain tissue and phantoms demonstrates substantially improved stability and robustness of local optic axis estimation. This work enables reliable depth-resolved fiber orientation mapping in sophisticated fiber crossing scenarios and extends the potential of single-input PSOCT for analyzing structural connectivity at high depth resolution.

## 1. Introduction

Polarization-sensitive optical coherence tomography (PSOCT) is an emerging modality for imaging fibrous biological tissues [1–5], offering depth-resolved microstructure information beyond that of intensity-based optical coherence tomography. By measuring the changes in polarization state of light as it propagates through tissue, PSOCT quantifies intrinsic optical properties such as birefringence and optic axis [6]. In *ex-vivo* brain imaging, these properties are particularly valuable for mapping the orientation of myelinated fiber tracts. Single input PSOCT is the most commonly used setup because of its speed and simplicity. Unlike full Mueller or Jones matrix approaches [7,8], which require complex hardware to modulate multiple input polarization states, single-input PSOCT is simple to implement and does not require extensive computational resources. With circularly polarized light incident on the sample, one measurement enables calculation of retardance and optic axis orientation. However, the apparent retardance and optic axis that single input PSOCT measures are a cumulative signal, affected by all the birefringent layers along the optical path [9]. Therefore, accurate recovery of the depth-resolved local optic axis is critical for resolving the true fiber orientation in depth, especially in brain imaging where it can enable structural connectivity reconstruction at mesoscopic to microscopic resolution.

Accurate local optic axis estimation under noise remains challenging, particularly for single-input systems. Previous methods, such as recursive Jones matrix inversion, are sensitive to noise, as depth-resolved local phase retardation calculation amplifies speckle noise [10], which becomes more severe in deeper tissue regions. Alternative approaches such as polarization state tracing are also sensitive to noise, as they rely on circle fitting [11] or differential vector estimation [12] on the Poincaré sphere in addition to reversing the cumulative polarization effect from previous birefringent layers. Circle fitting can also be illconditioned in low-birefringence regions, affecting the stability of local optic axis estimation [11,13].

To address these limitations, we developed a robust framework for local optic axis estimation in single-input PSOCT. We propose a novel geometric approach that integrates Polarization Super Pixels (PSP) [13] with a mirror state constraint on the Poincaré sphere [14]. The PSP method functions as a spatially adaptive binning scheme for tracing polarization change as light propagates, which allows for dynamically allocating larger kernels to low-birefringence regions to prevent mathematical singularities, while preserving resolution in high-birefringence areas. Furthermore, we apply a physical constraint based on the mirror state of the input polarization incident on the sample as any fitted circle defined by the local optic axis of a birefringent tissue will pass through the mirror state [14]. Multi-input PSOCT configurations inherently satisfy the physical boundary conditions of the mirror state due to their full polarization state coverage. In contrast, standard single-input PSOCT systems measure only a subset of the full Jones or Mueller matrix elements, leading to a loss of key polarization information. Here, we demonstrate that explicitly enforcing the mirror state constraint within a geometric framework compensates for this missing information, enabling robust local optic axis recovery with a single-input configuration. By reinforcing the fitted polarization trace to intersect the mirror state, we effectively constrain the geometric fit. This constraint significantly stabilizes optic axis estimation in low-SNR or deep tissue regions. This is particularly critical for low-birefringence areas, where the Stokes vectors fail to complete a full trajectory around the Poincaré sphere and thus remain far from the mirror state. We demonstrate the algorithm’s capability to recover stable local optic axis orientations in a bilayer phantom and biological tissue using a single-input PSOCT system, validating its effectiveness for robust fiber orientation mapping under noise.

## 2. Methods

### 2.1 The problem of cumulative retardance and optic axis

Polarization sensitive optical coherence tomography (PSOCT) measures the cumulative polarization signal as light propagates inside the sample. Conventionally, the input state is selected to be circularly polarized light to avoid overlapping with the sample’s optic axis on the Poincaré sphere [6,15]. The measured PSOCT signal, including the apparent retardance and apparent optic axis, can be simulated by modeling the local optic axis at every depth of the sample, treated as a pure retarder.

A simple simulated case consists of homogenous layers of constant local optic axis, for which the measured apparent retardance and optic axis are markedly different from the local ground truth (Fig. 1a, dashed line). For the retardance plots, the dashed line represents the cumulative retardance, and its slope corresponds to the local birefringence. When the apparent retardance accumulates to 90 degrees, two artifacts appear: retardance folding and optic axis flipping. First, the measured retardance begins decreasing instead of increasing and then reverses again after reaching 0, producing a characteristic folding pattern. Second, the apparent optic axis shifts by 90 degrees. On the Poincaré sphere, Stokes vectors trace a great circle that intersects the input circular polarization state (0,0,1) and its corresponding mirror state (0,0,-1). Each time the measured polarization state passes through these points, retardance folding and optic axis flipping occurs. The retardance is measured as the shortest arc length (geodesic distance) from the input polarization state to the moving Stokes vector representing the polarization state. As a result, the Stokes vectors trace the other side of the Poincaré sphere and retardance starts decreasing after passing through (0,0,-1). For the apparent optic axis, the Q and U components flip signs after passing through (0,0,-1), which adds 180 degrees to the azimuthal angle in the Q-U plane. Since the apparent optic axis is half of the angle between the Q and U components, this produces a 90 degree shift. This combined phenomenon is often referred as “banding” in previous literature [16,17].

**Fig. 1.**
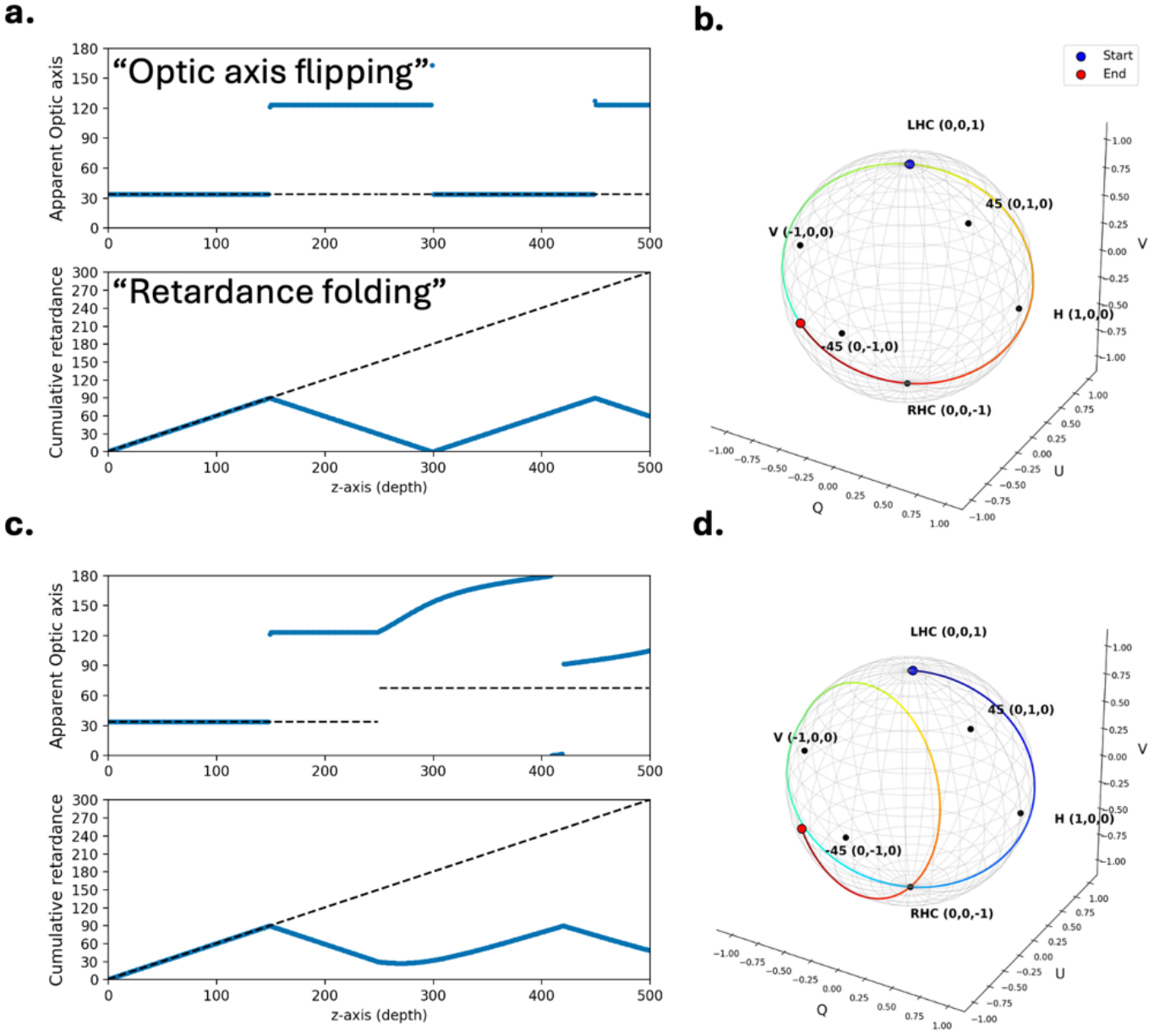
Illustration of measurement artifacts with single-input PSOCT in single-layer and multiple-layer fibers. (a) Simulated measurements of apparent optic axis (top) and cumulative retardance (bottom) with a constant optic axis, exhibiting two characteristic artifacts: optic axis flipping, where the recovered axis abruptly jumps by 90°, and retardance folding, where the cumulative retardance oscillates rather than increasing monotonically with depth, and (c) Simulated measurements of apparent optic axis (top) and cumulative retardance (bottom) with the multi-layer fibers with differed optic axis. (b) and (d) Corresponding trajectory of the polarization state on the Poincaré sphere with respect to (a) and (c).

When there are two or more different local optic axes along the imaging depth, the behavior of the measured apparent retardance and optic axis becomes more complex (Fig. 1c). The Stokes vectors initially trace a great circle but transition to tracing another circle when the second layer begins with a different optic axis orientation. Importantly, as shown in Fig. 1d, no matter how many layers there are of varying optic axis orientations and regardless of sample complexity, the Stokes vectors always trace a circle that passes through the mirror state, (0,0,-1) [14]. This forms the physical basis for the mirror state constraint described further in Section 2.2.

The aim of this paper is to recover the true local optic axis and local birefringence of the sample from the noisy measurements of apparent retardance and optic axis acquired with single input PSOCT.

### 2.2 Recovering local optic axis for multi-layer samples

The recovery of the local optic axis utilizes two mainstream approaches: recursive Jones matrix inverse and geometric polarization state tracing.

The recursive matrix inverse approach [9,18,19] models the layers as retarders. To obtain the local Jones matrix at depth N, the algorithm multiplies the measured cumulative matrix at that depth by the inverse of all the local Jones matrices above it:

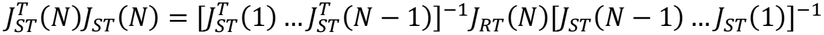

This operation removes the cumulative polarization effects of the overlying tissue, leaving only the local contribution. While conceptually straightforward, this method is highly sensitive to noise, as errors at superficial depths propagate and amplify with increasing depth in the sample.

The geometric approach determines the local optic axis by analyzing the trajectory of polarization states on the Poincaré sphere [11]. As light propagates through a birefringent sample, the measured Stokes vectors trace a circle on the sphere’s surface, who’s axis of rotation defines the apparent optic axis. In practice, this axis is commonly estimated by either fitting a plane to a cluster of Stokes vectors within a small sliding window or by using discrete differential geometry [12], where the normal vector to the fitted plane corresponds to the apparent optic axis. The local optic axis is then recovered by recursively inverting the rotations associated with the preceding layers. This method can be highly sensitive to noise depending on how the rotation axis is estimated, and it struggles when the sample’s birefringence is too weak for fitting a well-conditioned circle on the Poincaré sphere and estimation near layer interfaces is challenging [11]. The discrete differential geometric rotation axis approximation can be noise sensitive as well.

Local optic axis estimation is especially challenging in complex biological samples with varying optic axis orientations because low signal-to-noise ratio and low birefringence results in error propagation and instability in parameter fitting. We propose a constrained geometric method combining mathematical and physical aspects to address this problem: polarization super-pixels and a physical constraint based on the polarization mirror state.

We incorporate polarization super-pixels (PSP) as a preprocessing step which groups pixels with similar polarization states into clusters, from which only the centroids are used for downstream analysis [13]. In the case of single input PSOCT, it is necessary to impose spatial constraints to ensure that the pixels within a cluster are spatially connected in depth. PSP increases the SNR of the measured data while decreasing the effective number of layers processed during recursive matrix inversion, significantly improving the quality of the local optic axis estimation. PSP functions similarly to an adaptive binning scheme, where low-birefringence regions require more pixels and high-birefringence regions fewer pixels, mitigating the ill-conditioning issues associated with low-birefringence regions in the geometric method. Fig. 3 shows the apparent retardance and optic axis under noise, where the PSP centroids and cluster indices illustrate the adaptive nature of the method.

To obtain PSP, we adaptively cluster the pixels of polarization depth profile using a one-dimensional Simple Linear Iterative Clustering (SLIC) superpixel framework via scikit-image. Although SLIC natively operates on multidimensional image coordinates, we isolated the z axis by formatting the input array into an array of shape (*N*, 3), where *N* denotes the total number of pixels along depth. The three channels are the normalized polarimetric parameters: normalized phase retardation *δ*/(*π*/2), and the normalized trigonometric vector components of orientation (sin(θ) + 1)/2and (cos(θ) + 1)/2. The number of PSP is the key parameter of the SLIC algorithm, and in practice, we find that each PSP should contain around 10 pixels, so for 200 pixels imaging window in depth the number of PSP should be around 20.

The compactness parameter in the SLIC framework controls the balance between polarimetric feature homogeneity and spatial grid uniformity. High compactness forces spatial proximity to dominate the optimization metric, prioritizing uniform segmentation where PSP are driven to have nearly identical physical lengths regardless of underlying feature variations. Conversely, lowering compactness allows polarimetric feature similarity to dictate cluster boundaries. The compactness parameter can be tuned depending on the data, and a compactness of 20 was found to be a reasonable starting point for this work. Spatial connectivity is strictly enforced via standard connected-component post-processing along the single coordinate axis, guaranteeing that all generated PSP form continuous, non-overlapping 1D clusters.

We also enforce the physical constraint of the mirror state during local optic axis estimation. The mirror state is the polarization state on the Poincaré sphere that has the same linear components as the input light state but reversed helicity (circular component), meaning that in single-input PSOCT, a right-circular input state (0,0,1) has a mirror state of left-circular (0,0,-1). The mirror state is different for other PSOCT configurations. The recursive matrix inversion method implicitly enforces the mirror state constraint by modeling each layer as a linear retarder. Existing geometric methods calculate osculating planes using exclusively localized data subsets or consecutive moving vectors along the tracked depth profile, without anchoring the optimization space to a fixed global reference such as the mirror state. However, a previous study [14] shows that the circle traced by the Stokes vectors should always pass through the mirror state in PSOCT measurements, as shown in Fig. 2. This is a constraint that is explicitly enforced in the current work, when fitting circles using the geometric approach. Constraining the evolution to pass through the mirror state stabilizes circle fitting and can significantly enhance accuracy, particularly when if the Stokes vectors are far from the mirror state.

**Fig. 2.**
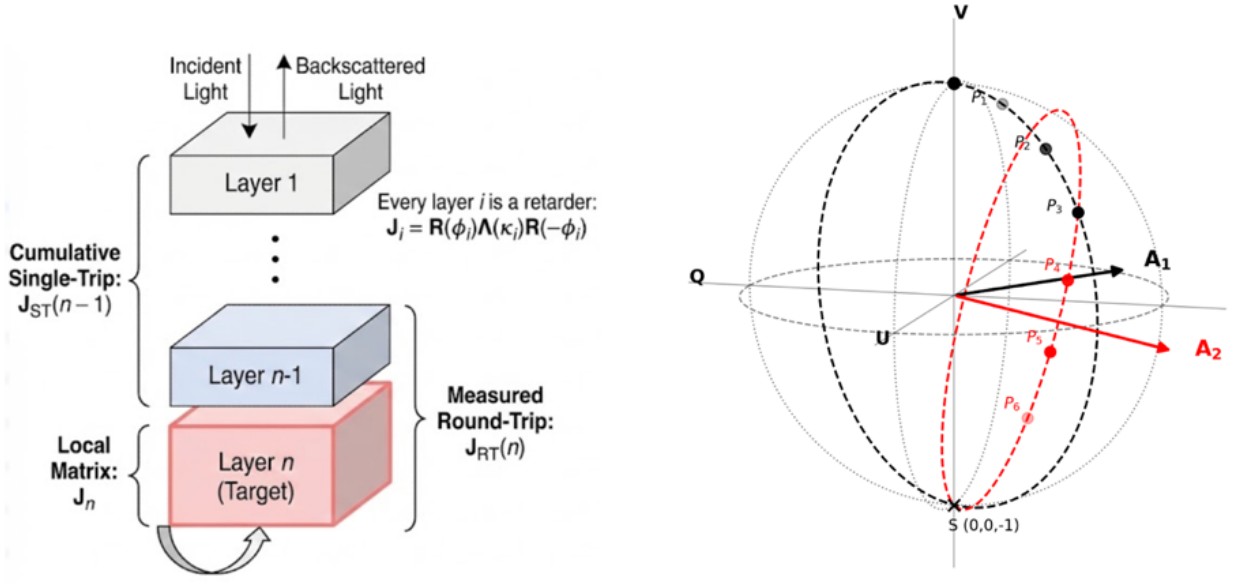
Illustration of existing methods for local optic axis estimation in PSOCT. (Left) Layered model used in the matrix inverse method, and (Right) Polarization state tracing on the Poincaré sphere, where the local optic axis is estimated by fitting the polarization trajectory to a great-circle arc.

**Fig. 3.**
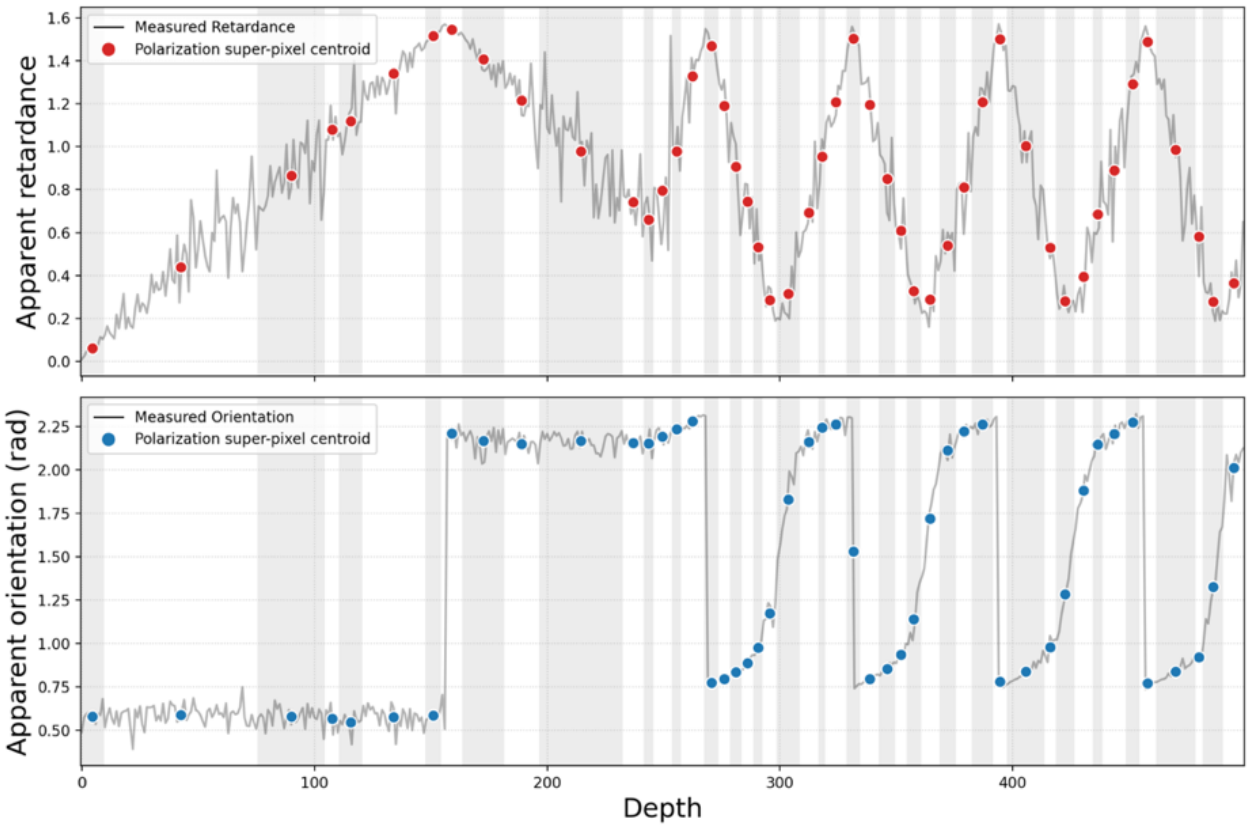
Polarization super-pixel (PSP) applied to PSOCT measurements in depth. Apparent retardance (top) and optic axis orientation (bottom) as a function of depth, with the raw measured signal shown in line and the PSP centroids overlaid as colored dots.

The complete constrained geometric method combines the recursive inverse method, polarization state tracing, polarization super-pixels, and the mirror state constraint. The measured apparent optic axis and retardance are first used to obtain the polarization super-pixels. The proposed algorithm works recursively through a layer-by-layer process, with each layer being defined as a small window of data points. Beginning with the first layer, a circle is fitted on the Poincaré sphere via Singular Value Decomposition (SVD) while explicitly incorporating the mirror state constraint. The local optic axis is extracted as the normal direction of the fitted circle. Using the estimated local optic axis, the local birefringence is optimized by minimizing the difference between the predicted and measured Stokes vectors in the layer. A Jones matrix is then constructed using the estimated local optic axis and birefringence, and its inverse is applied to all subsequent measurements to eliminate the cumulative polarization effects. Geometrically, this operation corresponds to rotating the measured trace backward on the Poincaré sphere, to reverse the polarization state to its original circular state. This procedure repeated for the next layer, treating it as the new initial layer until all depth-resolved polarization properties are retrieved.

The mirror state constraint is explicitly imposed by translating the data points by the mirror state, and then applying SVD. A 3D plane is defined by a normal vector 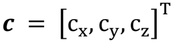 and a scalar distance offset *d*. Any point **p= [***x, y, z*]^T^lying on this plane must satisfy the linear equation:

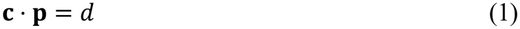

If we constrain our plane to pass through the mirror state **S= [**0,0, −1]^T^, then substituting **S**into the plane equation gives:

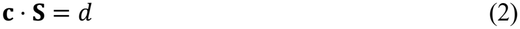

Subtracting these two equations eliminates the unknown offset *d* and leaves the equation:

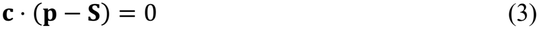

We now define a new shifted vector ***V*** = (**p** − **S**) and solve for **C**by applying SVD, essentially calculating ***V***^***T***^***V*** and calculating the eigendecomposition. The resulting fitted plane passes exactly through the mirror state, while the SVD yields the normal vector that minimizes the sum of squared orthogonal distances from the measured Stokes vectors to the plane. This produces the total least-squares optimal fit subject to the mirror state constraint.

### 2.3 Imaging setup and samples

PSOCT imaging was performed using a custom polarization-sensitive system based on a Telesto spectral domain PSOCT platform (Thorlabs, Newton, NJ, USA). The system operates at a 1300 nm central wavelength, with an axial resolution of approximately 2.5 µm in tissue and a lateral resolution of approximately 5 µm. A single input of circular polarization state was incident on the tissue sample.

To validate the local optic axis algorithm, we imaged two different samples: a birefringent phantom and a human brainstem sample. We first imaged a two-layer polycarbonate phantom with distinct optic-axis orientations fabricated following the method described by Liu et al. [1]. The phantom exploits the photo-elastic effect, in which birefringence is induced by stretching a polymer sample after heating it above its glass-transition temperature, producing samples that stably exhibit homogeneous birefringence. The stretched segments are assembled into phantoms with multiple well-defined layers of distinct birefringence and known optic axis orientations, making them well-suited for quantitative validation of PSOCT reconstruction methods. We next imaged a human brainstem sample provided by the CONNECTS program [20]. Human brain tissue was obtained from the Massachusetts General Hospital (MGH) Autopsy Suite. Samples were excised within a 24-hour post-mortem interval and fixed by immersion in 10% formalin. This study was approved by the MGH Institutional Review Board. The brainstem sample was refractive index matched using glycerol before and after agarose embedding and then imaged in mineral oil to maximize the depth penetration and signal to noise ratio.

## 3. Results

### 3.1 Theoretical simulation

To first validate the proposed method and compare it to the existing methods, we constructed a simulated dataset with controlled noise levels. The dataset contains three layers with different birefringence, each with a distinct local optic axis. To simulate measurement noise, we added Gaussian noise to the measured complex Jones (horizontal and vertical polarization channels), and then derived the measured retardance and orientation.

Fig. 4 shows the local optic axis and birefringence reconstruction result with different levels of noise. all three methods performed reasonably well at the low noise level (15%), successfully recovering the true local optic axis and retardance indicated by the red line. The same polarization super-pixel preprocessing procedure is applied to all methods. At the medium noise level (30%), the polarization state tracing (PST) method begins to fail while the recursive matrix inverse method and the proposed constrained geometric method both perform reasonably well in local optic axis estimation. When the noise is high and approaches realistic experimental noise levels (50%), only the constrained geometric method remains robust enough to reconstruct the true local optic axis accurately.

**Fig. 4.**
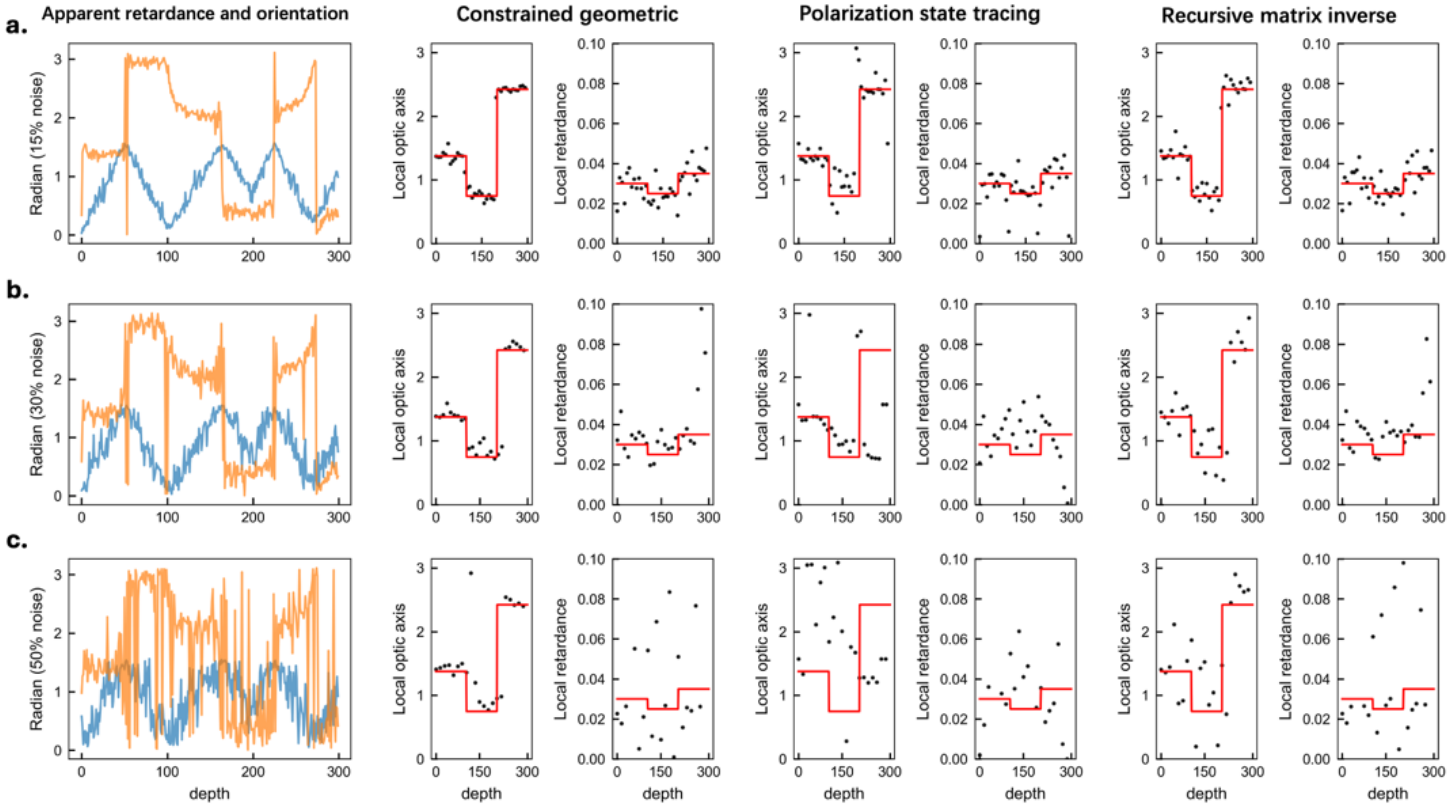
Comparison between simulation results of local optic axis estimation under increasing noise levels. Under each noise level, simulated apparent retardance and orientation (leftmost column) are shown alongside the recovered local optic axis and local birefringence from three methods: proposed constrained geometric, polarization state tracing, and recursive matrix inverse. Results are shown for (a) 15% noise, (b) 30% noise, and (c) 50% noise. The red line indicates the ground truth.

### 3.2 Phantom local optic axis estimation

In this experiment, two birefringent layers were stacked with a relative angle of 65° between their optic axes, as illustrated in Fig. 5a and in the average intensity projection of PSOCT measurements in Fig. 5b. The entire stacked phantom sample was then rotated through 180° to acquire measurements at 30°intervals. The local optic axis was recovered at each rotation position to validate the constrained geometric method’s ability to resolve distinct layers under rotation.

**Fig. 5.**
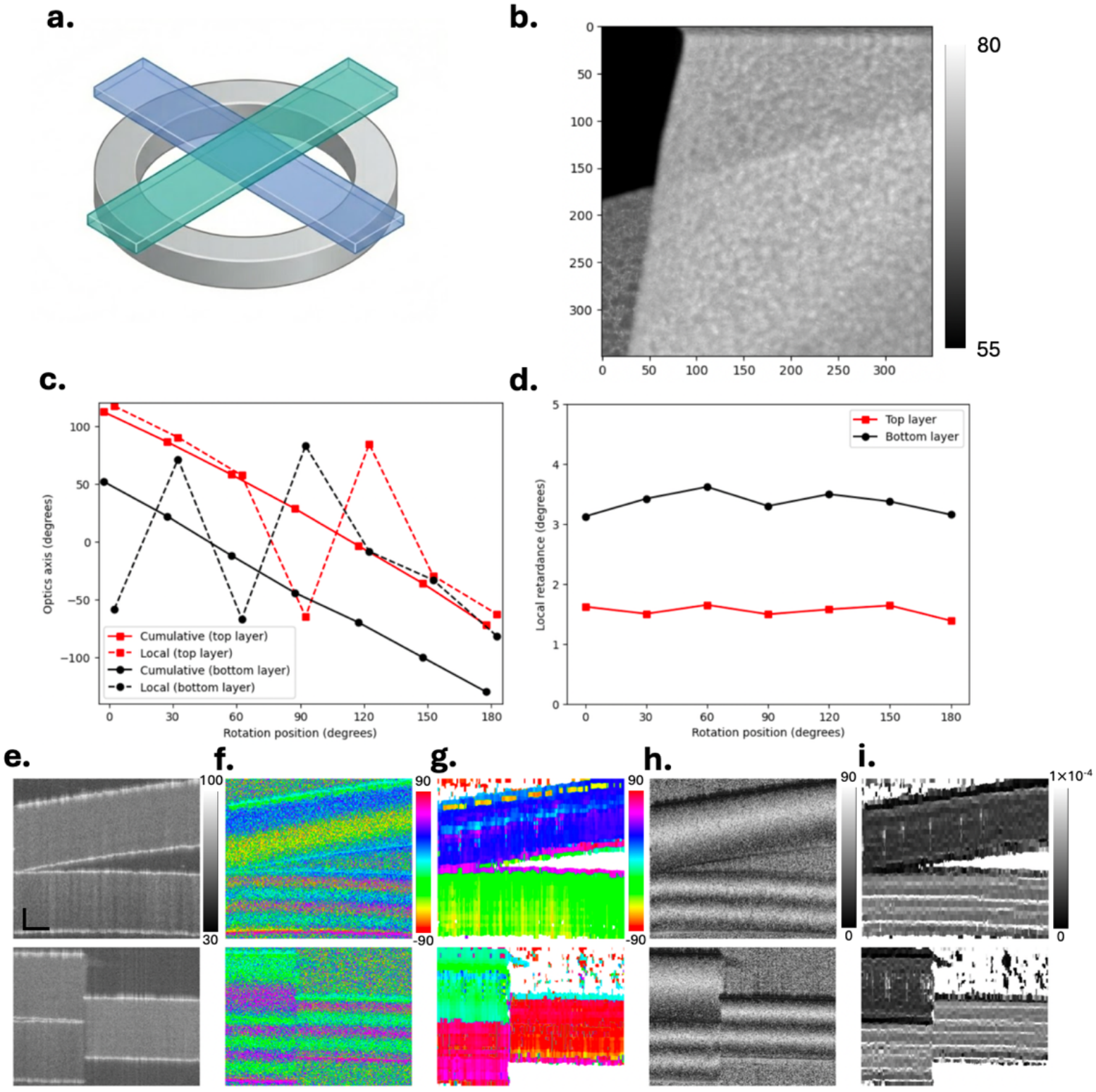
Validation of local optic axis estimation on a two-layer birefringent phantom. (a) Schematic of the crossed-layer phantom. (b) Representative average intensity projection showing the overlapping layers. (c) Recovered local optic axis orientation (solid line) as a function of phantom rotation angle for the top (red) and bottom (black) layers, along with apparent optic axis (dashed line). (d) Local birefringence of the top and bottom layers across rotation positions. (e–i) Representative cross-sectional images of intensity, apparent optic axis, local optic axis, apparent retardance, and local birefringence, respectively. (Scale bar: 100µ*m*for vertical and 500µ*m*for horizontal)

After reconstructing the local optic axis for the two layers, their mean values at each rotation angles are plotted, as shown in Fig. 5c. The solid line shows the local optic axis for the top layer (red) and bottom layer (black), respectively. It shows a clear linear trend for both the top and the bottom layers, with an average 64° angle between them. The dashed lines show the apparent optic axis measurements, which are difficult to interpret. The local birefringence indicates that the bottom layer is substantially more birefringent than the top layer (Fig. 5d). Fig. 5e-i show the cross-sections of intensity, apparent optic axis, local optic axis, apparent retardance, and local birefringence of the phantom sample. This method applies to other angles between sample layers, as demonstrated in the bottom row where the angle between layers is 90 degrees. The local optic axes in the two phantom layers are mostly uniform, with deviations near the interface between the top and bottom layers.

### 3.3 Brainstem local optic axis estimation

To assess the method’s performance in a realistic setting, we applied the proposed method to an ex vivo human brainstem sample.

Fig. 6 demonstrates that the local optic axis accurately resolves fiber orientation in the presence of crossing fiber tracts in a brainstem sample. Fig. 6a shows the cross-sections (indicated by the dashed white lines in Fig. 6b) of the brainstem sample. While the intensity is relatively uniform, the polarization contrast of the optic axis reveals intricate fiber organizations. Local optic axis estimation resolves varied fiber bundles crossing in depth: a superficial tract oriented at approximately +45°(indicated by the black arrow), and a deeper tract oriented at approximately −10° (indicated by the white arrow), that transitions beneath a fiber bundle oriented at −80° on the right. Despite this complex organization, the local optic axis recovers the orientation of the deeper fiber tract beneath the bundle above it. Fig. 6b presents the apparent and local optic axis maps at four depths. The apparent optic axis is corrupted by the overlying fibers, showing an erroneous orientation of ∼+30° where the true fiber direction is ∼−10°, indicated by white arrow in Fig 6b. The local optic axis, in contrast, resolves this difference in orientation, demonstrating its ability to disambiguate fiber crossings and recover depth-resolved optic axis information that the apparent optic axis cannot.

**Fig. 6.**
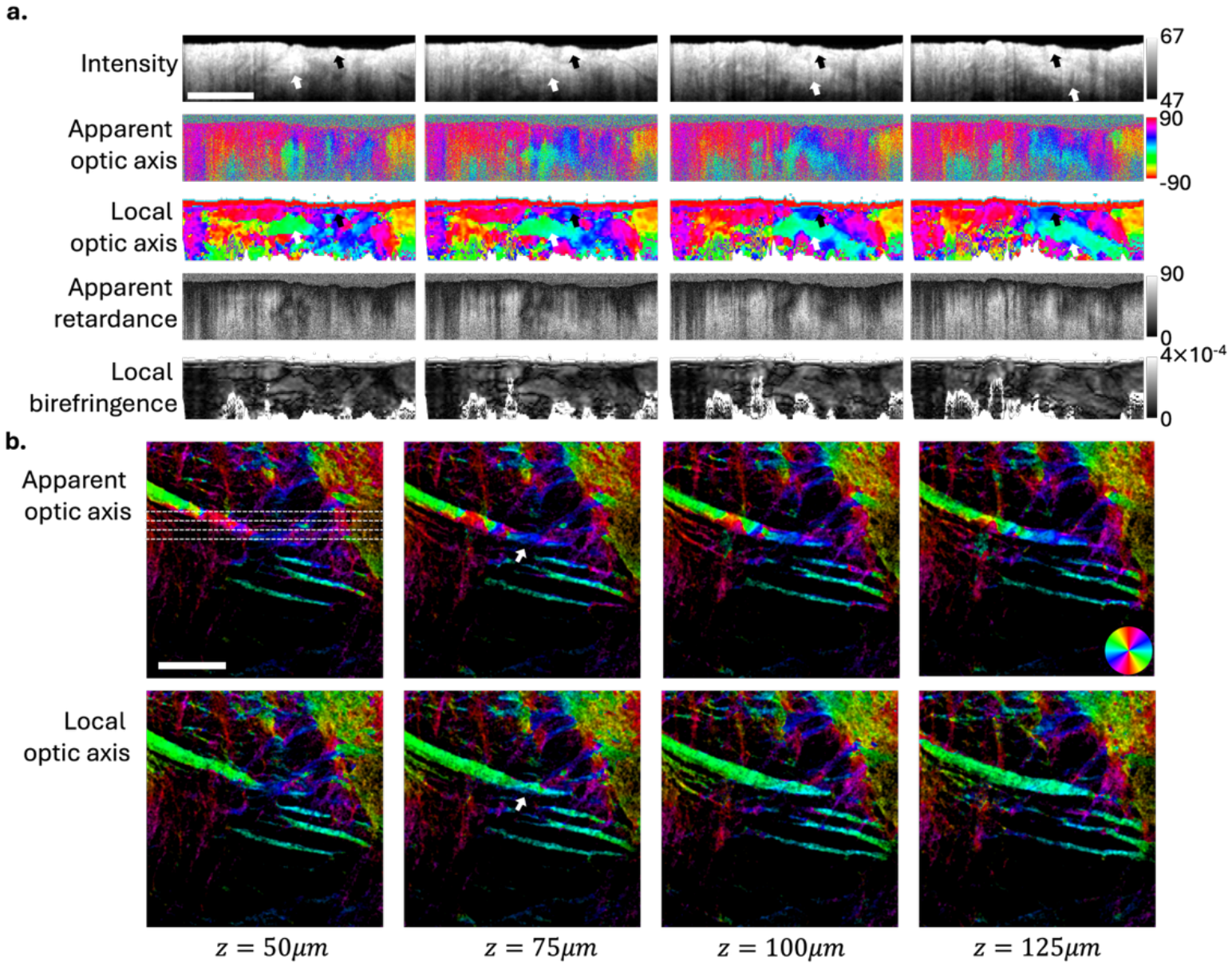
PSOCT imaging of an ex vivo human brainstem sample. (a) Cross-sectional images at four locations showing, from top to bottom: intensity, apparent optic axis, local optic axis, apparent retardance, and local birefringence. Black and white arrows in the intensity images indicate two distinct fiber tracts traversing in depth. (b) En face views of apparent optic axis (top) and local optic axis (bottom) at depths z = 50, 75, 100, and 125 µm.

## 4. Discussion

We have proposed a robust method for local optic axis estimation under high-noise conditions in single-input PSOCT systems by combining polarization super pixels, a physical constraint based on the mirror state, and a Stokes-based geometric fit. Theoretical simulations, birefringent phantoms, and ex vivo human brain tissue samples were used to validate the method. PSP enables adaptive binning of pixels in depth, where low-birefringence regions use more pixels to mitigate the ill-conditioning and errors in the geometric fit, trading spatial resolution for improved signal-to-noise ratio in polarization features. It also adaptively adjusts the cluster size, becoming finer around thin fibers and coarser around thicker ones, preserving the fine structural information. Constraining the polarization state tracing to pass through the mirror state stabilizes the circle fitting and significantly enhances accuracy, particularly when Stokes vectors are distant from the mirror state. It is worth noting that under ideal, noise-free conditions, enforcing the mirror-state constraint on geometric tracing introduces no additional physical information compared to the recursive matrix formulation, and both approaches inherently converge to the same solution. However, PSOCT measurements bear significant speckle noise. Using a mirror-state constraint under a geometric framework effectively transforms the mathematical matrix inversion chain into a geometric path-tracing problem. This formulation provides an implicit regularizing mechanism, making the reconstruction more robust against the noise in the measurement and numerical instabilities.

The local optic axis enables two broader applications in volumetric brain mapping. The first is depth-resolved reconstruction of the in-plane fiber orientation throughout the imaging volume, which provides significantly improved axial resolution in the orientation map derived from PSOCT. Without local optic axis estimation, even in homogeneous samples, there is a fundamental trade-off: higher intrinsic birefringence improves the polarization signal-to-noise ratio but amplifies the banding artifacts. This effectively restricts usable information to shallow depths. Building on the local optic axis, the method has the potential to enable full 3D axis orientation estimation at high resolution. Current 3D axis orientation approaches rely on enface birefringence and orientation maps, without leveraging the depth-resolved capability of PSOCT measurements. Since the local optic axis provides depth-resolved orientation at the scale of tens of microns, incorporating it into 3D axis orientation reconstruction could increase the effective depth resolution by an order of magnitude and enable 3D fiber tracking.

We acknowledge that the constrained geometric method has several limitations. Using PSP assumes that the underlying biological structure is locally homogeneous within the scale of the super-pixel window. Therefore instead of resolving the sample in depth at its intrinsic axial resolution, the effective resolution varies depending on the birefringence strength and signal to noise ratio. Hyperparameter selection, such as the number of polarization super-pixels, the compactness, and the number of points used during circle fitting may require fine-tuning depending on the complexity of the sample. The geometric constraint relies on the mirror state and when the measurements lie close to the mirror state, the mirror state provides little useful information to improve the quality of the fit. This challenge could be addressed by a recent method that uses spectral polarization dispersion to multiplex multiple polarization measurements into a single acquisition [21].

## Data availability

The data that support the findings of this study are available from the corresponding author upon reasonable request.

